# A survey of duckweed species in Southern Italy provided first distribution records of the hybrid *Lemna* × *mediterranea* in nature

**DOI:** 10.1101/2024.08.09.607168

**Authors:** Leone Ermes Romano, Luca Braglia, Maria Adelaide Iannelli, Yuri Lee, Silvia Gianì, Floriana Gavazzi, Laura Morello

## Abstract

Interspecific hybridisation and polyploidization are two main driving forces in plant evolution, shaping genomes and favouring evolutionary novelty and ecological adaptation. Recent studies have demonstrated hybridisation within the genus *Lemna* (Lemnaceae Martinov) as well as triploid. *L*. × *mediterranea*, a recently described hybrids between *Lemna minor* and *Lemna gibba* was identified among only long lasting germplasm collections of *in vitro* propagated plants, originally collected at different times in the Mediterranean area.

We report the first distribution record of *L. × mediterranea* in the nature, in the Campania region of Southern Italy, the same area where *Lemna symmeter* was described as a new species about 50 years ago, confirming their synonymy. Eight specimens isolated from five different sampling sites over an area of about 4200 Km^2^ showed identical genetic profiles by Tubulin-Based Polymorphism (TBP) analysis, suggesting their common origin from the same hybridisation event, followed by clonal dispersal. The *L. × mediterranea* population of Campania is genetically different from any of the previously analysed clones, suggesting that recurrent hybridisation between the parental species may occur. The natural hybrid clone is triploid, with *L*. *gibba* as the plastid donor, and remarkably similar to it by morphology, although the typical gibbosity of this species becomes evident only upon *in vitro* flower induction. Flowers are protogynous and self-sterile. Ecological factors including competition with parental and invasive species, niche and climate change adaptation, stability in time and space likely played a role in the successful establishment of *L. × mediterranea*.

**Highlights:**

- Interspecific hybridisation within the genus Lemna documented in nature
- β-tubulin intron length polymorphism for tracking duckweed species distribution
- Flower induction in Lemna × mediterranea

## 1. Introduction

Lemnaceae Martinov is a family of floating aquatic plants that populate lentic and slow-moving water bodies, thriving, particularly, in systems impacted by cultural eutrophication (Akinnawo, 2023). The family accounts for 35 species and two hybrids (Appenroth et al., 2024) a number still subject to updating and revision, thanks to novel genetic and genomic approaches. Duckweeds are often addressed as the most miniature flowering plants (Landolt, 1986). Although most species can produce fully fertile flowers, fruits, and tiny seeds, Lemnaceae prefer vegetative propagation as a reproductive strategy. They are the fastest-growing angiosperms (Ziegler et al., 2015) in fact, some species can double their biomass in less than two days (Zaoh et al., 2014). Their exceptionally short lifespan, rapid asexual growth and straightforward *in vitro* cultivation have made plants from the genus *Lemna* L. attractive as a model organism for fundamental ecological and evolutionary research (Laird and Barks, 2018; Acosta et al., 2021). In the late nineteenth century, Darwin described how species adapted and evolved under the influence of natural selection. Research has thoroughly investigated the effect of natural selection on population genetics and evolution, but the matter is still far from a complete understanding (Rieseberg, 2001; Abbott and Brennan, 2014). The introduction of modern molecular techniques to population studies has been a critical factor in unravelling the natural adaptation of species to different environments and their interaction with one another (Rieseberg and Willis, 2007). An important opportunity for experimenting entirely new genetic combinations is represented by interspecific hybridization, as has been underlined numerous times in recent years (Wong et al., 2022). This process can be defined as the gametic interaction across taxonomically distinct species, resulting in the formation of novel organisms that may eventually lead to hybrid speciation (Hörandl, 2022). Both plant geneticists and crop breeders have recognized the importance of intra- and interspecific hybridisation as tools for crop improvement and selection. Species hybridisation and introgression occur naturally and play a crucial role in plant adaptation to the natural environment (Rieseberg and Willis, 2007), being essential parts of plant speciation (Rieseberg and Willis, 2007; Warner and Walworth, 2010). Five distinct genera (*Spirodela, Lemna, Landoltia, Wolffia* and *Wolffiella*) represent the Lemnaceae family, occupying important ecological niches (Bog et al., 2019). Their floating nature sees them as a regulatory factor of the biodiversity co-existing in the water column underneath (Romano and Aronne, 2021; Feller et al., 2024). Their fast growth, adaptation to various sources and concentrations of nutrients (Fang et al., 2007) and tolerance to different environmental conditions (Ziegler et al, 2023; Romano et al., 2024), are fundamental factors influencing their coexistence with other floating plants in the natural environment. The efficient reproductive strategy of these tiny plants confers them an advantage over other competitors and increases the risk of eutrophication in the wetlands (Feller et al., 2024), making them good biological indicators of eutrophication conditions. Due to the growth promoting effect of rising environmental temperatures, the duration of duckweed dominance in slow moving freshwaters of temperate Europe is increasing (Peeters et al., 2013). More so, faster-growing alien species are spreading and, in some cases, competing with autochthonous ones (Ceschin et al., 2016). Various research has shown that modelling their growth patterns is essential to estimate the potential increase in their abundance and mitigate their effect on the natural environment (Peeters et al., 2013; Feller et al., 2024). The combination of nuclear molecular markers and plastid DNA barcoding, allow a more precise taxonomic categorization of these plants, thus obviating the limited number of morphological traits. Genomic approaches make these plants amenable to population studies, biogeographical investigation and cross-species comparative work, facilitating modern molecular ecology and evolutionary biology research.

As for other higher plants, interspecific hybridisation also occurs within the Lemnaceae family as it has been recently demonstrated (Braglia et al., 2021a, b). The application of the TBP (Tubulin Based Polymorphism) molecular marker has been a critical factor in re-categorizing the species *Lemna japonica* Landolt, as an interspecific hybrid between *L. minor* and *L. turionifera* (Braglia et al., 2021a). Another interspecific hybrid, between *L. minor* L. and *L*. *gibba* L., referred to as *L*. × *mediterranea* Braglia and Morello, has been identified in the framework of the genetic characterization of an *ex-situ* germplasm repository (Braglia et al., 2024), held at the CNR - Institute of Agricultural Biology and Biotechnology (Morello et al., 2024). Due to the geographic origin of the investigated accessions, this hybrid was supposed to correspond to a putative new species identified in the Campania region (Italy) and described in 1973 as *Lemna symmeter* by Giuseppe Giuga, closely related to *L. gibba,* but differing from it in flower development and sterility (Giuga, 1973). The aims of this paper were: i) to retrieve wild populations of *L*. *× mediterranea* in the natural environment in the same region where *L. symmeter* was originally described, ii) to verify the possibility that the two taxa are the same iii) to provide information about distribution and biodiversity of the hybrid and its parent species. Interspecific hybrids, coming from plants endemic to a particular geographical area, are an unprecedented source of information. They adapt well, compete with parent as well as alien species, can be recollected at different time intervals, and are used as biomonitoring in studying the novel adaptation of plant species to a particular geographical area. Furthermore, sterile hybrids among facultative sexual species, producing pure clonally propagated lineages with identical genetic background, make it easier to investigate their somatic mutation rate, lifespan and frequency, and can help scientists to better understand the effect of induced environmental changes at a fast vegetative reproductive cycle (Mo et al., 2022; Tao et al., 2022; Du et al., 2022).

## 2. Materials and methods

### 2.1 Plant habitat identification and in-field sampling

To investigate the presence of the nothotaxon *L*. × *mediterranea* in Italy and verify its identity with the putative species *L*. *symmeter*, as described by Giuga in a monographic leaflet (Giuga, 1973), we have planned and conducted extensive plant sampling campaigns, conducted during June, July, and August of 2022 in the Campania region. This season was chosen due to the presence of the optimal conditions for duckweed growth: higher ambient temperatures and reduced water volumes.

To plan the site visits, we adopted a two-split approach: (1) gathering historical data from Giuga’s documentation and (2) analysing satellite images using Google Earth Pro®. To validate the satellite imagery data, we implemented the following steps:

1. Waterway maps consultation: we studied detailed maps to identify all possible waterways in the study area.
2. Classification of waterways: the waterways were then classified into types, as per certain specific definitions that applied to the scope of our work, based on their size, flow rate, and surrounding vegetation.
3. Vegetation presence assessment: employing the ‘historical imagery’ tool in Google Earth® (https://earth.google.com/web), for each waterway, we investigated the presence of green vegetation over time. This step was used as an additional verification step and was not used as exclusion factor.
4. Extraction of non-matching watercourses: at this stage, we excluded watercourses that did not reflect consistent vegetation presence or did not meet the criteria set at higher steps.

This helped us ensure that sites selected from the analyses of satellite images matched actual plant habitats identified through historical records.

The waterways have been classified into three main categories: Artificial Waterways (AW), a category for all the manmade flowing waterbodies (canals and drainage channels); Natural Waterways (NW), acronym used for all the waterways represented in the category of streams, rivers, creeks, gullies, springs, or washes; Artificial Ponds (AP), including steady waters wells, artificial lakes and water reservoirs (see Table 1). This integrative approach enhanced the precision of the habitat identification process.

**Table 1.** Fieldwork sampling summary where the column Sampling group shows the different samples collected during the expeditions; the column Accession describes the ID assigned to each specimen after morphological analysis; the column TBP categorizes the species to which each clone belongs; the column Type of waterways describes the different waterways: Artificial Waterways (AW), Natural waterways (NW), Artificial Ponds (AP); the column Coordinates indicates the geographical position of the different samplings sites.

| Sampling Group | Accession TBP |  | Type of waterway | Coordinates |  |
| --- | --- | --- | --- | --- | --- |
|  |  |  |  | N | E |
| S1 | LER001 | *Lemna minuta* | AW | 40°57.3113' | 14°01.7699' |
| S2 | LER002 | *Lemna gibba* | AW | 40°58.1384' | 14°01.4786' |
| S2 | LER003 | *L. minuta* | AW | 40°58.1384' | 14°01.4786' |
| S3 | LER004 | *Wolffia arrhiza* | AW | 41°00.8751' | 14°00.8985' |
| S3 | LER005 | *L. gibba* | AW | 41°00.8751' | 14°00.8985' |
| S3 | LER006 | *L. minuta* | AW | 41°00.8751' | 14°00.8985' |
| S3 | LER007 | *L. gibba* | AW | 41°00.8751' | 14°00.8985' |
| S3 | LER008 | *L. minuta* | AW | 41°00.8751' | 14°00.8985' |
| S3 | LER009 | *L. gibba* | AW | 41°00.8751' | 14°00.8985' |
| S4 | LER010 | *L. gibba* | NW | 41°02.7979' | 14°03.1177' |
| S4 | LER011 | *L. gibba* | NW | 41°02.7979' | 14°03.1177' |
| S5 | LER012 | *L. minor* | AW | 41°00.0776' | 14°17.3984' |
| S6 | LER013 | *L. minuta* | AW | 41°00.0403' | 14°20.1625' |
| S6 | LER014 | *Lemna × mediterranea* | AW | 41°00.0403' | 14°20.1625' |
| S7 | LER015 | *Lemna minor* | AW | 40°58.3651' | 14°27.4156' |
| S8 | LER016 | *L. gibba* | AW | 41°00.021' | 13°59.160' |
| S9 | LER017 | *L. minor* | AW | 41°15.7129' | 14°52.9815' |
| S10 | LER018 | *L. minor* | AW | 41°00.481' | 13°58.518' |
| S10 | LER019 | *W. arrhiza* | AW | 41°00.481' | 13°58.518' |
| S11 | LER020 | *L. × mediterranea* | AW | 40°45.0092' | 14°31.6233' |
| S11 | LER021 | *L. × mediterranea* | AW | 40°45.0092' | 14°31.6233' |
| S11 | LER022 | *L. × mediterranea* | AW | 40°45.0092' | 14°31.6233' |
| S11 | LER023 | *n. d.* | AW | 40°45.0092' | 14°31.6233' |
| S11 | LER024 | *W. arrhiza* | AW | 40°45.0092' | 14°31.6233' |
| S12 | LER025 | *L. × mediterranea* | AW | 40°30.9748' | 15°05.0356' |
| S12 | LER026 | *L. minor* | AW | 40°30.9748' | 15°05.0356' |
| S12 | LER027 | *L. × mediterranea* | AW | 40°30.9748' | 15°05.0356' |
| S13 | LER028 | *L. minor* | AW | 40°30.2460' | 15°05.5225' |
| S13 | LER029 | *L. minor* | AW | 40°30.2460' | 15°05.5225' |
| S14 | LER030 | *L. minor* | AP | 40°25.8012' | 15°13.0328' |
| S15 | LER031 | *L. minor* | NW | 40°21.9662' | 15°35.7301' |
| S16 | LER032 | *L. trisulca* | NW | 40°21.3626' | 15°36.8687' |
| S16 | LER033 | *L. minuta* | NW | 40°21.3626' | 15°36.8687' |
| S16 | LER034 | *L. minor* | NW | 40°21.3626' | 15°36.8687' |
| S17 | LER035 | *L. minuta* | NW | 40°21.1460' | 15°37.9503' |
| S17 | LER036 | *L. minor* | NW | 40°21.1460' | 15°37.9503' |
| S18 | LER037 | *L. minor* | NW | 40°21.2062' | 15°38.1312' |
| S19 | LER038 | *L. minor* | NW | 41°26.2255' | 14°13,2147' |
| S20 | LER039 | *L. × mediterranea* | NW | 41°24.3377' | 14°06,3202' |
| S21 | LER040 | *L. minuta* | NW | 41°25.1195' | 14°05,5152' |
| S21 | LER041 | *L. gibba* | NW | 41°25.1195' | 14°05,5152' |
| S21 | LER042 | *L. × mediterranea* | NW | 41°25.1195' | 14°05,5152' |

During the field expedition we visited approximately 130 different sites (**Figure S1**). When the presence of duckweeds in the waterway was verified, we performed a thorough examination to ensure a comprehensive collection of a representative sample.

As waterways were not always accessible, we employed a set of tools to sample plants in some cases. We specifically utilized telescopic nets for open water areas and buckets attached to long ropes for reaching otherwise inaccessible points. We collected samples manually in locations where direct access to the waterways was possible (Figure 1).

**Figure 1.**
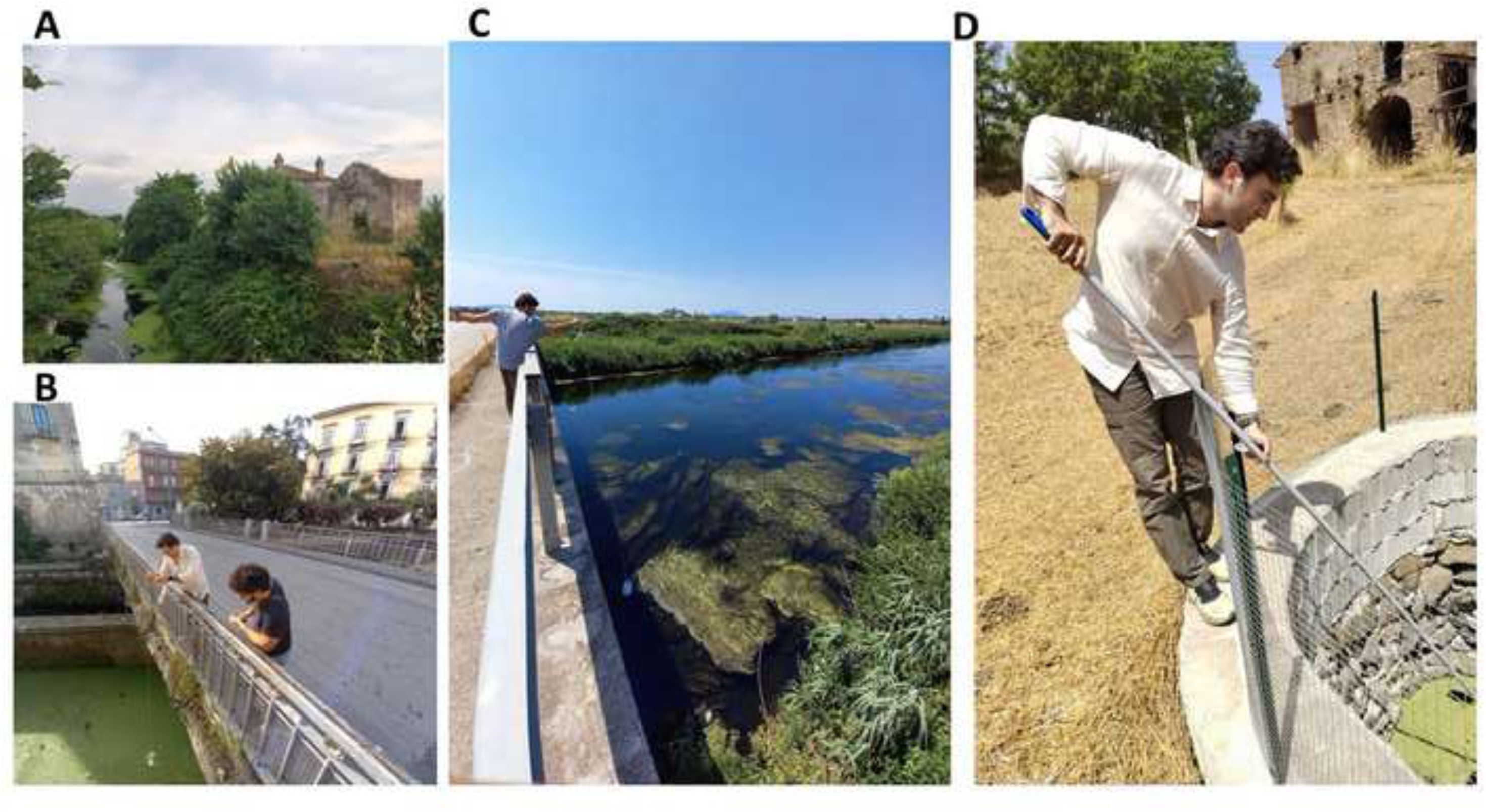
Different water bodies and collection strategies. Representative aquatic environments present in the Campania region, according to the reported categorization and indication of the sampling site. A) A canal at the historical site of Real Sito della Lanciolla (AW; S6); B) The canal crossing the Pagani city centre (AW; S11); C) The Volturno river (NW; S3); D) A water well in the National Park of Cilento (AP; S14).

Each plant sample gathered during sampling was catalogued by the sampling group (Table 1) to which georeferenced coordinates were assigned (Garmin Fenix 6 watch). Upon return to the laboratory, collected plants were rinsed with fresh water to remove any adhering debris or contaminants. Subsequently, the plants were examined under a stereo microscope to assess their morphological traits. Although morphological trait analysis is not sufficient for reliably discriminating different species within the family Lemnaceae, we performed this analysis to separate the putatively different specimens. Following the morphological analysis, individual colonies were isolated for sub-culturing, to produce clonal progeny. Plant sub-culturing was performed in a controlled environment at 20°C, under 16/8 hours’ photoperiod in a thermostatic incubator (Velp Scientifica, Italy). Upon reaching sufficient biomass in the sub-cultured material, surface sterilization according to Appenroth’s protocol (2015) prepared the samples for subsequent axenic propagation and genetic analysis. Forty-two specimens have been clonally propagated and labelled LER001-042 by the name of the collector, according to the recommendations of the International Steering Committee on Duckweed Research and Applications (Lam et al., 2020), as summarized in Table 1.

### 2.2 Plant material

Clones *Lemna gibba* 7742a and *Lemna minior* 5500 were investigated for the morpho-physiological analysis as a part of the CNR - Institute of Agricultural Biology and Biotechnology (CNR-IBBA duckweed collection - https://scientific-collections.gbif.org/collection/24fdf8b9-2165-44a5-9e30-019a2190331d) germplasm repository, constantly propagated in axenic conditions.

### 2.3 DNA extraction

DNA was extracted by grinding 50-100 mg of frozen fronds, in Eppendorf tubes with 3 steel beads and a few mg of quartz sand with a TissueLyser II (Qiagen, Hilden, Germany) at 30 Hz for 90 sec. Plant tissue was lysed following the standard procedures of the DNeasy Plant Kit (Qiagen, Hilden, Germany). DNA was eluted in a final volume of 100 µL of 5mM Tris-HCl and stored at −20°C.

#### TBP amplification, capillary electrophoresis and data analysis

TBP amplification was performed according to the protocol reported by Braglia et al. (2020). Fluorescence labelled amplicons were separated by capillary electrophoresis on a 3500 Genetic Analyser (Thermo Fisher Scientific Inc.,) as described by Braglia et al. (2023). Each sample DNA was independently analysed twice. The amplicon sizing and allele detection of the TBP electropherogram gained by both intron regions (I and II intron) was performed by Gene Mapper Software v. 5.0 (Thermo Fisher Scientific Inc., Germany). TBP profiles were included in a local database together with all previously analysed accessions for subsequent analysis. A detailed list of the additional Lemna clones of which TBP profile were collected in the local database is reported in **Supplementary Table 1.** The peak size (base pairs) and height (RFUs) of each pherogram were collected through a Microsoft Office Excel file and all the TBP profiles were aligned according to the peak size.

Limited to the three related taxa *L. gibba*, *L.* × *mediterranea* and *L. minor*, the TBP peaks (markers) scoring was performed considering both intron regions, in order to estimate genetic similarity and a single presence/absence matrix (1/0) was then generated. The FAMD (Fingerprint Analysis with Missing Data) program, v.1.31 (Schlüter and Harris, 2006) was used to estimate genetic parameters: percentage of polymorphic markers, number of fixed markers, and number of private alleles (only meaningful if groups are mutually exclusive) found in each of the three analysed taxa. Multivariate analyses were inferred using Jaccard’s similarity index implemented in Past 4 software (v. 4.13) for Windows (Hammer et al., 2001) and UPGMA (Unweighted Pair Group Method with Arithmetic mean) trees and a principal component analysis (PCA) were constructed. The measure of how faithfully the designed dendrogram preserves the estimated genetic distances was evaluated through the Cophenetic correlation coefficient using the Past 4 software. The same software were also used to interrogate the TBP matrix in order to estimate: (dis-) similarity values by a pairwise comparison analysis as the proportion of the shared diversity to the total diversity (Whittaker, 1972) within and between the three *Lemna* taxa (Koleff et al. 2003); the Shannon’s diversity index, as the measure of the clone richness and relative abundance, within the three taxa.

### 2.4 Plastid marker analysis

The *atpF-atpH* spacer and *rps16* barcoding region were used as plastid sequence markers for *Lemna* sp. and *Wolffia* sp., respectively. Primers and PCR conditions were as reported by Braglia et al. (2021a) and Bog et al. (2013). Amplicons were purified with the Microclean kit (Labgene Scientific, Châtel-Saint-Deni, Switzerland) and sent to an external facility for sequencing on both strands (Microsynt, Balgach, Switzerland). After trimming and polishing, sequences were either aligned using the BioEdit alignment tool to identify SNPs or used as probes for BLAST analysis (https://blast.ncbi.nlm.nih.gov/Blast.cgi).

### 2.5 Frond Area

Among the different clones collected for the experiment, we selected three, LER002, LER012 and LER014, as representative of the hybrid and its parental species, for frond size comparison. More specifically, we expected *L.* × *mediterranea* to show significant differences in frond size when compared to the parent species *L. minor* and *L. gibba,* as reported by Braglia et al. (2021b, 2024). Plants cultivated under the same environmental parameters were imaged after 168h of growth. To image the plants, we have utilized an Olympus SZX9 stereo microscope, equipped with a Sony Alpha II camera. Thirty fronds per clone were measured utilizing the Image J software as described by Romano et al. (2022).

### 2.6 Flower induction

*Lemna gibba* (7742), *L*. × *mediterranea* (LER021) and *L*. *minor* (5500) colonies were grown on diluted (1:20) modified Hutner’s medium (Hutner, 1953) lacking NH_4_^+^ and supplemented with 1% sucrose, based on Kaihara et al. (1981) under long-day conditions (16:8 light/dark cycle) at 25°C and a fluorescence light intensity of 160 µmol/s/m^2^ in presence of 30 µM salicylic acid (SA). Fifteen *L.* × *mediterranea* fronds were fixed on 0.3% agar-added medium. Flower development was monitored through observations and photographs at time intervals of three to 15 hours, starting 11 days after flower-inducing culture initiation. For comparison, *L. minor* and *L. gibba* flower development was investigated, and images of mature flowers and pollen production were taken.

## 3. Results

### 3.1 Genetic identification of the isolated clones, geographic distribution, and plant associations

Upon visiting about 130 sites across an area of about 4200 km^2^, populations of Lemnaceae were found at 21 sites (Figure 2). As morphological identification in the Lemnaceae is no longer considered sufficient for species determination, all subcloned samples were genetically identified. Three *Wolffia* samples were easily identified by morphology at the genus level, but species assignment was confirmed by sequencing the *rps16* plastid marker. BLAST analysis of the three identical sequences obtained (https://blast.ncbi.nlm.nih.gov/Blast.cgi, accessed on 06/24/2024) retrieved *rps16* of *Wolffia arrhiza* (L.) Horkel ex Wimm. clone 8272 (ID HE819982.1) as the highest scoring based on best match, with a 98.55% identity over 968 nucleotides. The three *W. arrhiza* specimens shared identical TBP fingerprinting profiles, suggesting they belong to the same clonal population, closely related to other Italian clones included in the CNR-IBBA duckweed collection, but genetically more distant from other clones (unpublished data).

**Figure 2.**
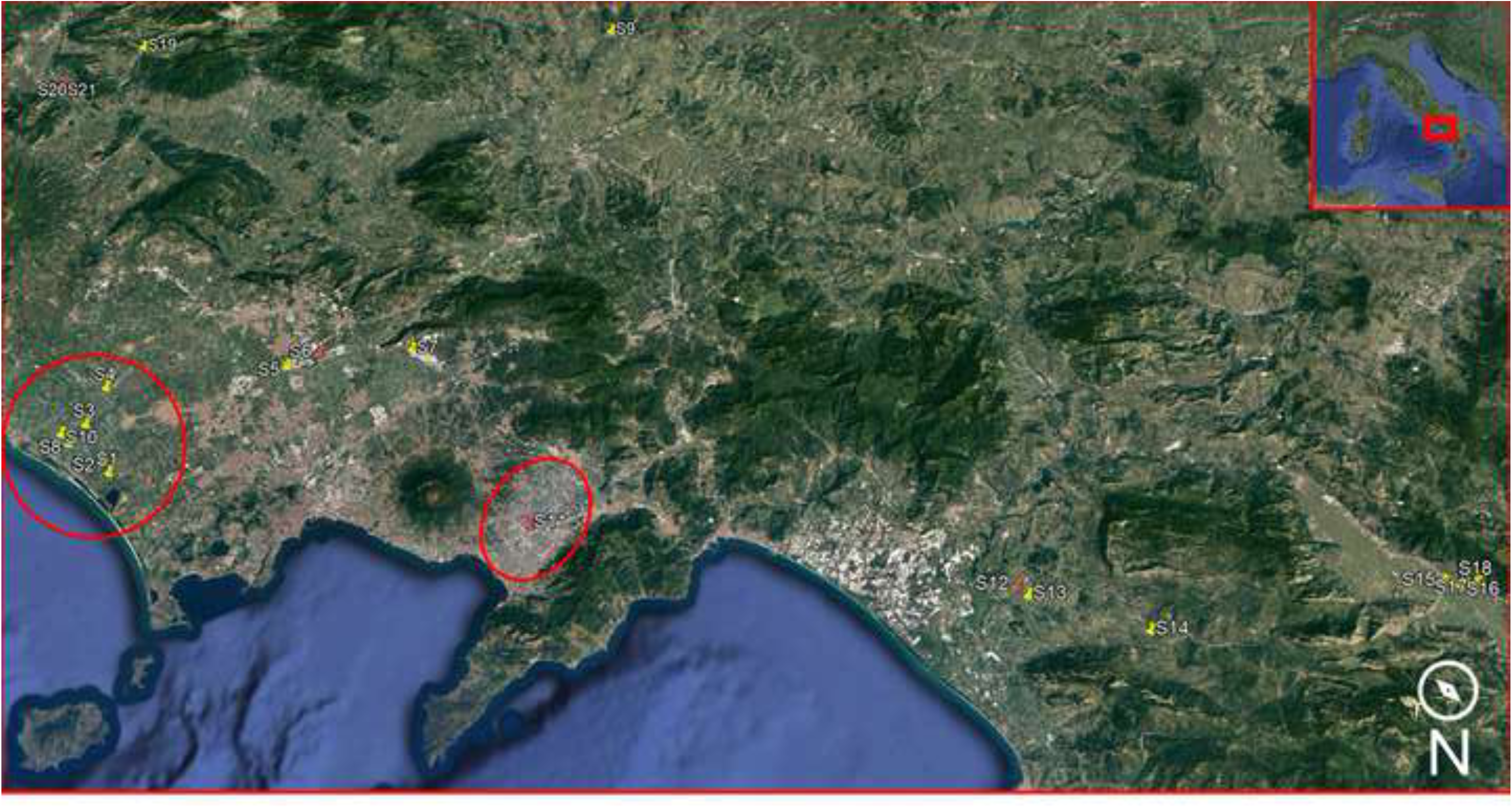
Map of the sampling campaign. Yellow and red pins indicate sampling sites; red pins indicate sampling sites in which there was also occurrence of *Lemna* × *mediterranea* (see Table 1). Red circles show the sites in which the putative species *L. symmeter* was described. Georeferenced sampling data are available at the following link: https://earth.google.com/earth/d/1UqwplzeEjHGrYZZjv_Haor_urxYyV3gW?usp=sharing

Each subcloned *Lemna* specimen was genetically identified exclusively by the TBP marker, able to discriminate all closely related *Lemna* species and their hybrids (Braglia et al., 2021a, b).

Table 1 reports the species identity of 41 duckweed clones (LER023 was lost before analysis), belonging to *W. arrhiza* and to four *Lemna* species: *L*. *gibba, L*. *minor, L*. *minuta* Kunth and *L*. *trisulca* L. In addition, we were also successful in retrieving in nature *L*. × *mediterranea*, the interspecific hybrid between the two autochthonous species *L*. *gibba* and *L*. *minor*, so far reported only from *ex-situ* germplasm collections (Braglia et al., 2021b, 2024). The geographic distribution of the 21 sampling sites is reported in Figure 2. *Lemna minor* was the most common species, found in 12 out of 21 sites, followed by the alien species *L. minuta*, at seven sites, *L. gibba, and L. × mediterranea,* at five sites each, and *W. arrhiza* recovered from three sites*. Lemna trisulca* was found at one site only. The presence of a single species occurred at ten sites, while association of two or three species were found at eleven sites.

### 3.2 Characterization of Lemna × mediterranea specimens

*Lemna* × *mediterranea* was identified from the nature for the first time in this work, in the Campania region, where *L*. *symmeter* was described as a new species (Giuga, 1973), and no more reported since then. Populations were found at five different sites (S6, S11, S12, S20 and S21) distributed over an area of at least 4200 km^2^, alone or in association with either *L. minor* or *L. gibba*, but in no case with both parental species. One of the sites, S6, actually falls within one of the two main areas described for *L. symmeter.* Unfortunately, none of the plants collected and described by Giuga were preserved as herbarium specimens and no molecular comparison with the clones collected in this study could be made.

All eight specimens collected, LER014, 020-022, 025, 027, 039 and 042, showed identical TBP profiles for both the first and second β-tubulin intron, suggesting they have a clonal origin from a single hybridisation event (see below).

Absolute genome size measurement of clone LER027 (758 Mbp) and comparison with previously investigated *L.* × *mediterranea* clones from the CNR-IBBA duckweed collection showed it was in the same size range of two accessions from South Tyrol, Italy (9248; 780 Mbp) and Northern Germany (9425a; 774 Mbp), respectively. These two clones were found to be triploid, with two *L. gibba* and one *L. minor* subgenome contributions, using a qPCR approach (Braglia et al., 2024). Other hybrids from the *ex-situ* collection were instead found to be homoploid. Sequencing of the *atpF-atpH* intergenic plastid region identified *L. gibba* as the maternal parent for the eight *L.* × *mediterranea* clones isolated in this work, as well as for the two triploid collection clones mentioned before. a*tpF-atpH* sequences were 100% identical to each other (not shown).

### 3.3 Frond morphology

Frond morphology showed closer similarity of hybrids to *L. gibba* than to *L. minor*, with a less elongated shape (Figure 3). According to Braglia et al. (2021b), morphological analysis revealed larger fronds in triploid hybrids when compared to the parental species. The comparison of frond areas revealed significant differences among the different taxa (p < 0.001). Specifically, the frond areas were as follows: LER012 (*L. minor*) had an area of 5.19 ± 2.02 mm², LER014 (*L.* × *mediterranea*) had an area of 7.46 ± 2.39 mm² and LER002 (*L. gibba)* had an area of 4.57 ± 1.68 mm², (Figure 3).

**Figure 3.**
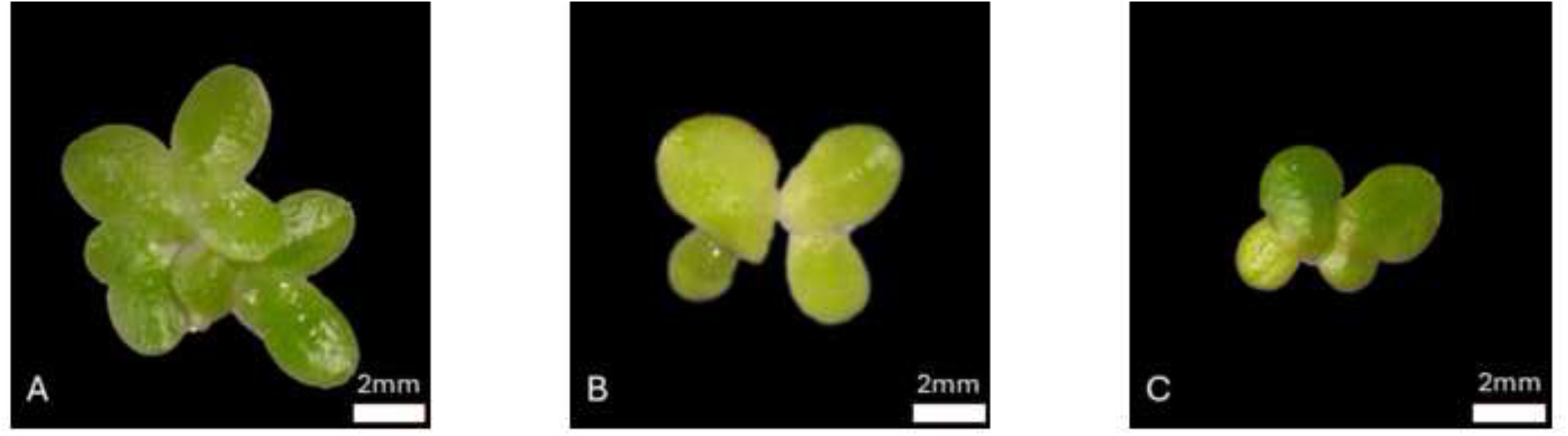
Representative images of frond colonies of the parental and hybrid taxa. A) *L. minor* LER012, B) *L.* × *mediterranea* LER014, and C) *L. gibba* LER002.

In addition, vein number was always five, like in *L. gibba*, when measured in ten fronds (**Supplementary Figure 2**). This number is typically three in *L. minor* and 3-4 in the diploid *L.* × *mediterranea* hybrid (Braglia et al., 2024).

### 3.4 Flower development

One of the most distinctive traits mentioned as critical for *L. symmeter* species determination in Giuga’s monograph, was flower development. In *L. gibba* the first anther appears together or soon after the pistil, later followed by the second anther (Landolt, 1986). Protogyny was instead characterized in *L. symmeter* by the first appearance of the stigma, followed some days later by the simultaneous growth of both stamens together, when the pistil was already withered. We then induced flowering *in vitro* in one *L.* × *mediterranea* clone (LER021) in parallel with *L. minor* 5500 and *L. gibba* 7742a, for comparison.

A first effect, starting after about seven days of SA treatment, was the increase in frond thickness in *L. gibba*, conferring it the classical gibbous morphology that was not observed under normal laboratory cultivation conditions. The same effect was reported for EDDHA treatment (De Lange and Pieterse, 1973). No such effect was seen in *L. minor,* while hybrids showed some degree of volume increase of the aerenchyma, although less pronounced *L. gibba* (Figure 4).

**Figure 4.**
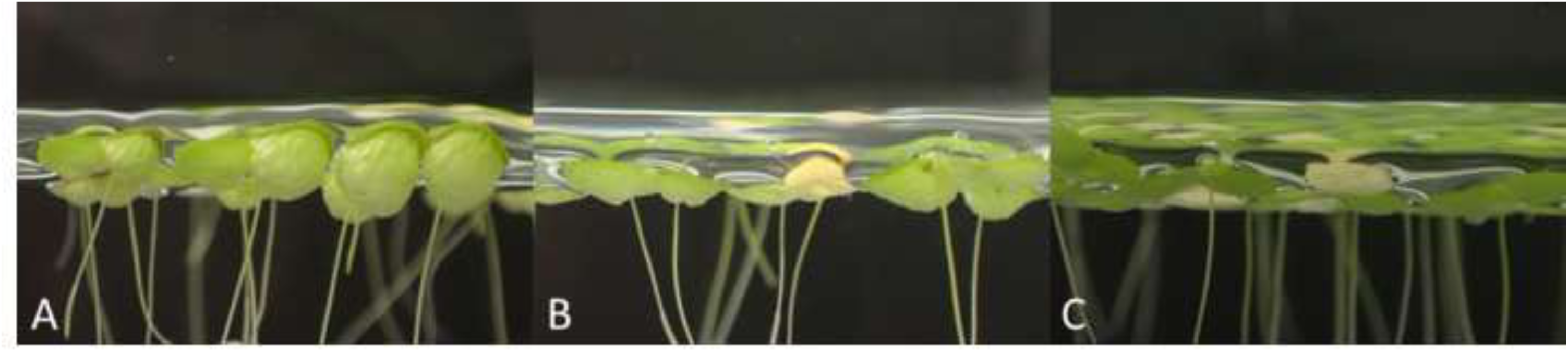
Frond morphology after SA addition, lateral view. A) *Lemna gibba* (7742a); B) *L*. × *mediterranea* (LER021); C) *L*. *minor* (5500). The photos were taken 19 days after flower induction. The hybrid *L.* × *mediterranea* has a visibly intermediate sized aerenchyma between *L*. *gibba* and *L*. *minor* when they flower under *in vitro* conditions.

Salicylic acid treatment at 30 μM successfully induced flower formation in *L.* × *mediterranea* (LER021), observed as early as day 11 (Figure 5A). A total of 17 flowers were monitored for six days. Due to the short and irregular duration of some organs, their monitoring could not be strictly scheduled. Notably, pistil emerged first, exhibiting prominent exudation on the stigma (Figure 5B). Following pistil maturation (Figure 5C), stamen growth initiated, frequently appearing yellow or pale yellow and morphologically similar to fertile stamens. Despite the initial wilting of the first stamen, the pistil remained viable in most cases, albeit with a reduced level of stigma exudate (Figure 5D). Deviations from the pathway described were also observed. In some cases, the first stamen developed after the pistil completely withered, in other cases it did not develop even several days after the pistil had already withered. Following the first stamen wilting, the second stamen emerged from the frond pouch and apparently matured, but anthers did not burst despite the apparently normal flower development (Figures 5E and F). Anther dehiscence was detected in less than 50% of the scrutinized stamens, as seen in Figure 6, while it was clearly visible in the parent species (Figure 7) as it was also reported for other *Lemna* species (Fourounjian et al., 2021, Lee et al., 2024). Figure 7 illustrates the floral structure of the parental species, *L. minor* (5500) and *L. gibba* (7742a) in the same inducing conditions. Notably, both species achieved complete anther maturation, evident by visible pollen release. As no fruit and seed setting was observed in LER021, further investigations are needed to assess if this is due to self-incompatibility or, more likely, to altered pollen viability and/or ovule fertility.

**Figure 5.**
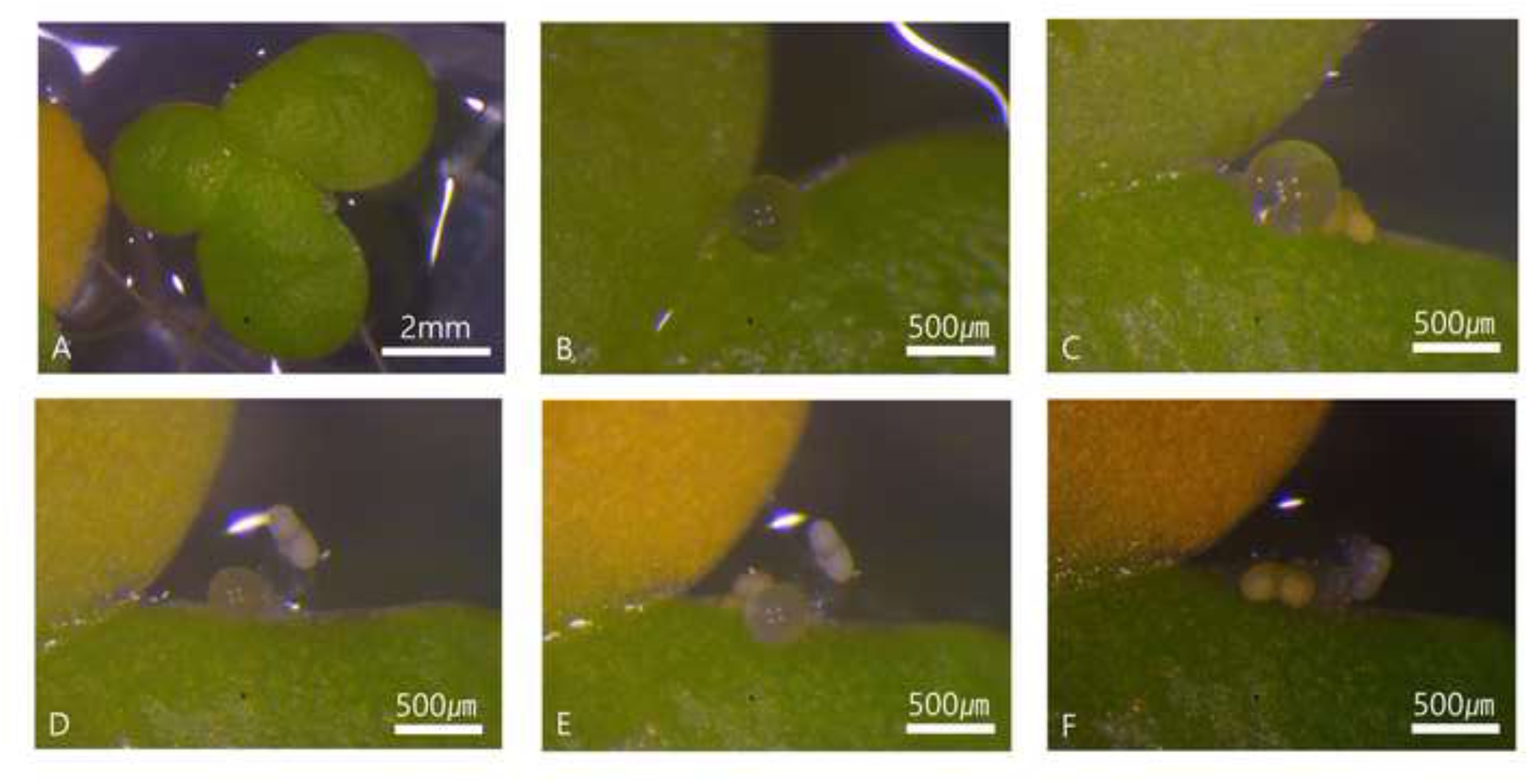
Flower development of *Lemna* × *mediterranea* (LER021). A) Whole flowering plant; B) Droplet on mature stigma; C) First stamen emergence and viable pistil; D) Pistil viability persistence despite wilting of the first stamen; E) Second stamen emergence; F) Fully developed second stamen following pistil and first stamen wilting.

**Figure 6.**
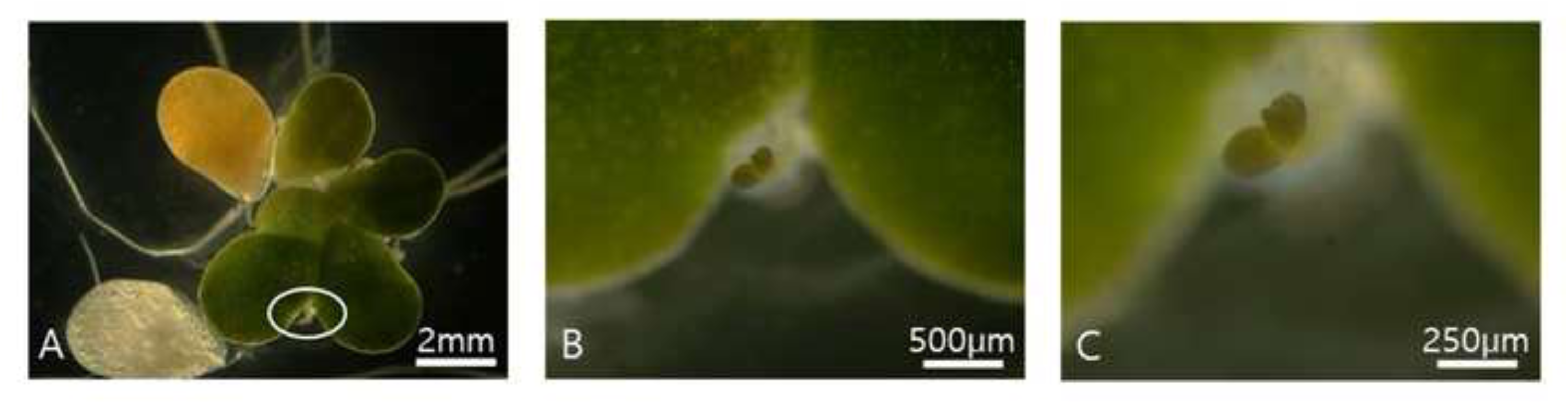
Flowering *Lemna* × *mediterranea* (LER021) plant A) Whole flowering plant; B) anthers opening for pollen release; C) magnification showing the vertical slit.

**Figure 7.**
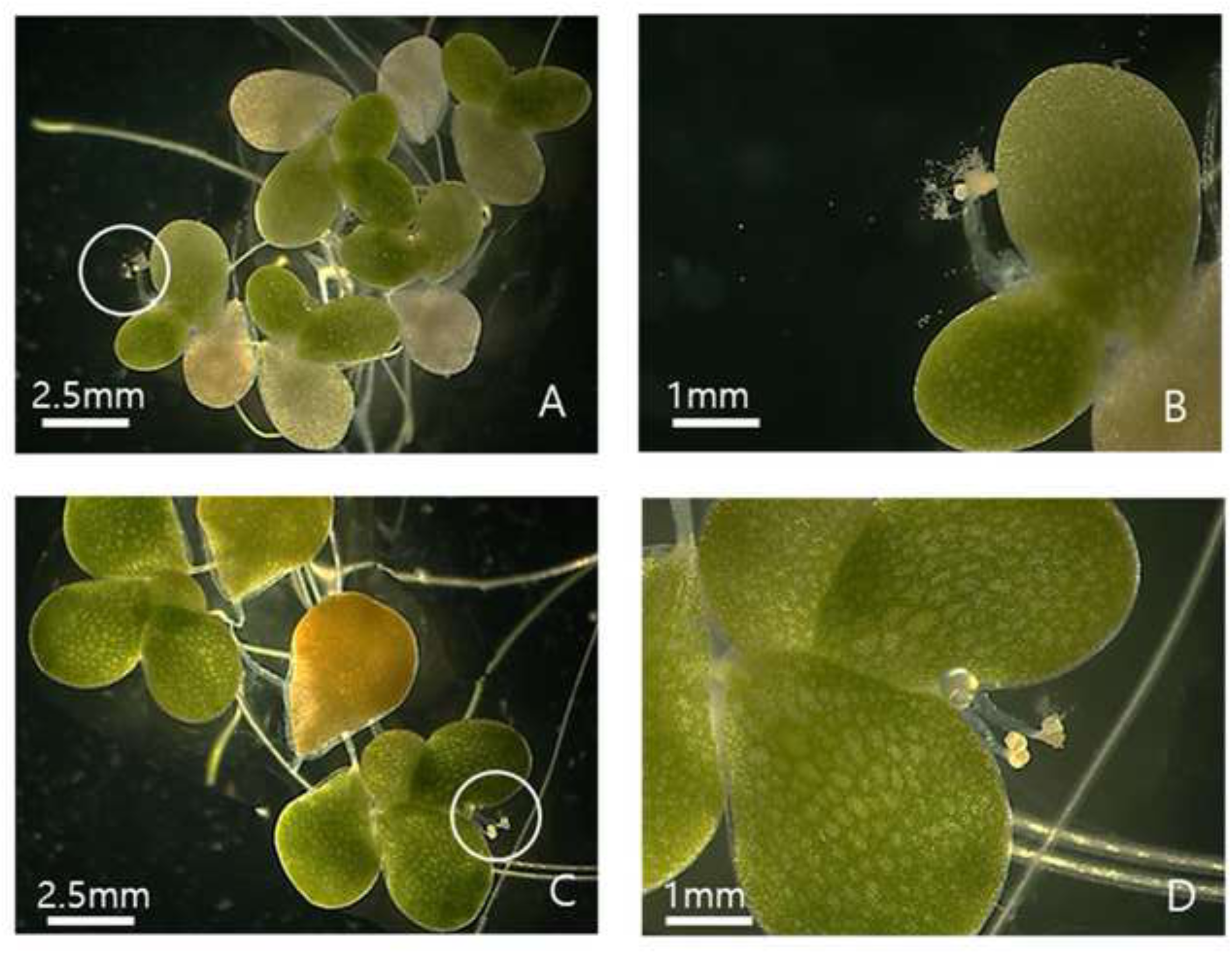
Flowers of *Lemna minor* (5500; A-B) and *L*. *gibba* (7742a; C-D). A-C) Whole flowering plants; B-D) Close-up view of flowers with dehiscent anthers releasing pollen.

### 3.5 Kinship and intraspecific genetic diversity in Lemna species

The peaks scoring of 29 TBP profiles, limited to *L. minor*, *L. gibba* and *L*. × *mediterranea*, revealed 37 markers, all polymorphic, 21 and 16 from the I and II intron region, respectively. Although TBP investigates a limited number of loci and has lower resolution than other markers in the low-variability species *Spirodela polyrhiza* (L.) Schleid. (Bog et al., 2022a), it was able to score genetic diversity in *L. minor* and *L. gibba* sample sets of the present paper. When each species was considered separately, a different rate of intraspecific genetic variation was highlighted by the marker scoring: the highest allelic polymorphism was recorded in *L. minor* (15 polymorphic and 11 private markers); *L. gibba* accessions revealed only five polymorphic and three private markers, while no variation was recorded within *L.* × *mediterranea*.

Furthermore, the genetic relationships between accessions of the three *Lemna* taxa are shown in the UPGMA dendrogram in Figure 8, obtained by cluster analysis of the genetic similarity estimated on the TBP data (cophenetic correlation coefficient 0.9678). As the variability between duplicate analysis was zero (not shown) and there was low risk to overestimate observed differences due to the limited resolution of the adopted marker, we considered each specimen as an independent clone when the Jaccard’s coefficient of similarity was below 1, with respect to any other. As already mentioned, all *L.* × *mediterranea* specimens were then identical among each other and clustered with *L. gibba*, with which they shared more alleles than with *L. minor.* The eight *L.* × *mediterranea* specimens in this study are then considered as a single clone and will be thereafter named LER-Lme. By applying the same rule, at least four different clones were found for *L. gibba* and 10 for *L. minor*. In this contest, the mean Jaccard’s similarity estimated within taxa showed a higher value in *L. gibba* (0.8769) than in *L. minor* (0.6690), further highlighting a greater genetic variability within this latter.

**Figure 8.**
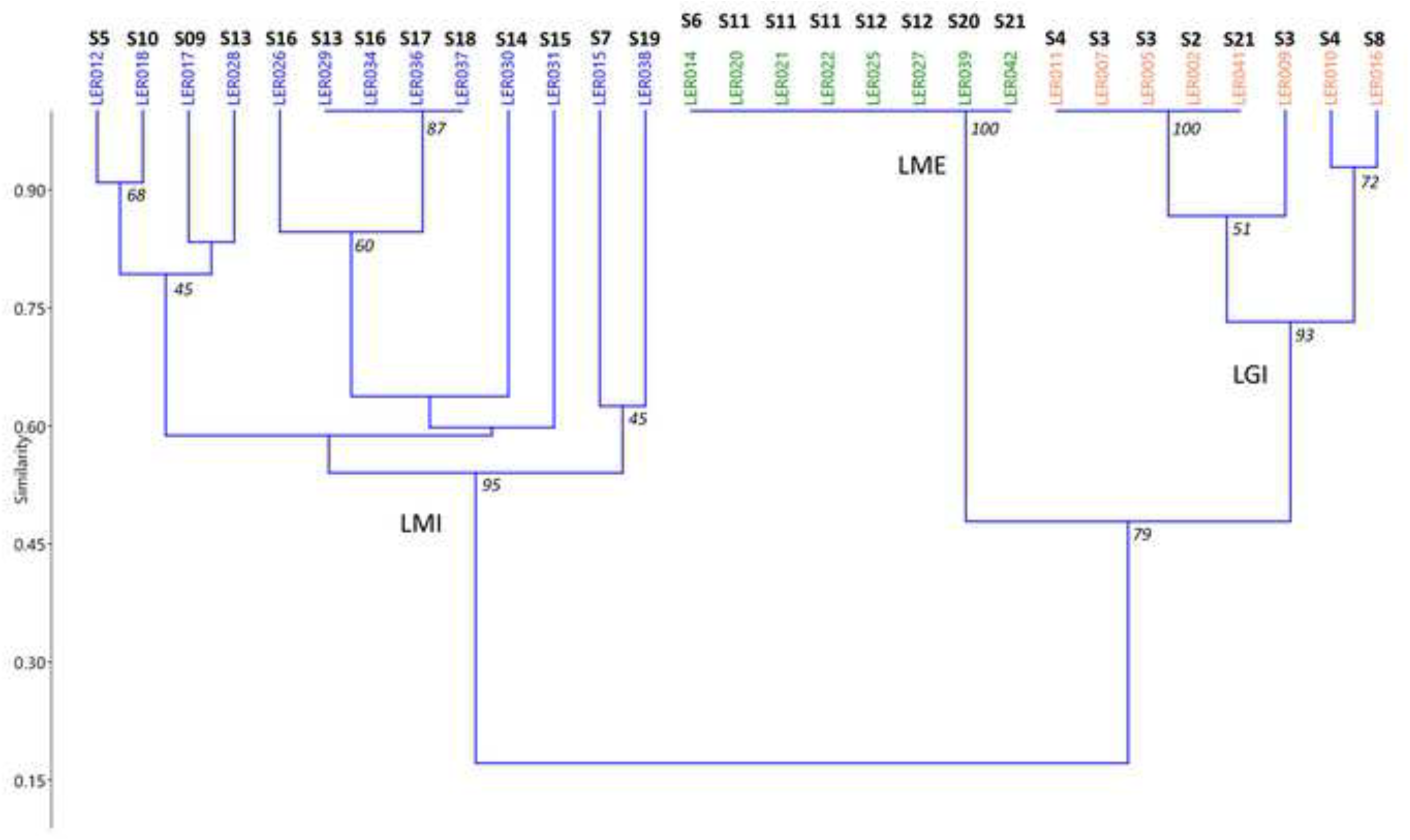
UPGMA dendrogram of the genetic distances among all collected *L*. × *mediterranea* clones and those of the parent species, estimated on the TBP data. Sampling sites are indicated for each sample. The acronyms LMI, LGI and LME refer to *Lemna minor, L. gibba* and *L*. × *mediterranea* respectively.

Specimens belonging to the same *L. minor* clone (LER029, LER034, LER036 and LER037) were found at four different sites (S13, S16-S18), while at one site, S13, specimens representing two distinct clones were collected (Figure 8). The same was also true for *L. gibba* (sites S3, S4), despite the lower variability scored by the marker in this species.

In addition, to identify putative parental clones of LER-Lme, a pairwise comparisons, according to the Whittaker formula, was estimated considering only those specimens classified as *L*. × *mediterranea* and its parental species (**Supplementary Table 2**). According to **Supplementary Table 2**, identity between accessions becomes evident as much as the value approaches 1. Conversely, dissimilarity approaches zero when specimens are indistinct, then considered as belonging to the same clone. As expected, the smallest mean value (0.00) was recorded comparing the eight *L.* × *mediterranea* specimens, while the highest value (1.00) was estimated between *L. minor* and *L. gibba* accessions. In this regard, single specimens from each *L. gibba* and *L. minor* group in this study (hereafter LER-Lgi and LER-Lmi) could be identified as putative parental clones involved in the hybrid formation as those showing the lowest dissimilarity values with respect to LER-Lme (**Supplementary Table 3**). When compared to *L.* × *mediterranea*, *L. minor* LER017 and *L. gibba* LER016 clones revealed the lowest dissimilarity values (0.39 and 0.28, respectively), and can be considered the most related to the hybrid. In agreement, the TBP pherogram comparison revealed a perfect allele overlap between these putative parental clones and the hybrid clones (**Supplementary Figure 3**).

A 3D PCA was inferred considering all *L. minor, L. gibba* and *L.* × *mediterranea* clones included in the CNR-IBBA duckweed collection. Despite the limited number of considered TBP markers, the 70% of the total variance was explained by the first three axes (48, 14 and 10% respectively, Figure 9) and all the analysed clones clustered into three distinct groups according to the three taxa. The LER-Lme hybrid clone (green diamond in Figure 9) was clearly distinct from all other LME clones. LER-Lmi (blue triangles in Figure 9) and LER-Lgi (yellow squares in Figure 9) clones did not form separate clusters, although they were mostly concentrated at one border of the respective species distribution cloud. Concerning *L. minor*, the most densely populated PCA cluster, the mean Shannon’s diversity index (SI) estimated within the group, showed non-significant difference comparing only LER-Lmi clones (SI = 2.92 ± 0.14) with all the clones of the CNR-IBBA duckweed collection (SI = 2.95 ± 0.13). This result remarks that, despite the limited number of analysed loci, the genetic diversity and the allelic richness characterizing the few collected clones of LER-Lmi do not differ to those estimated for a more numerous and geographically widespread germplasm resource represented by clones included in the IBBA Duckweed dataset duckweed collection.

**Figure 9.**
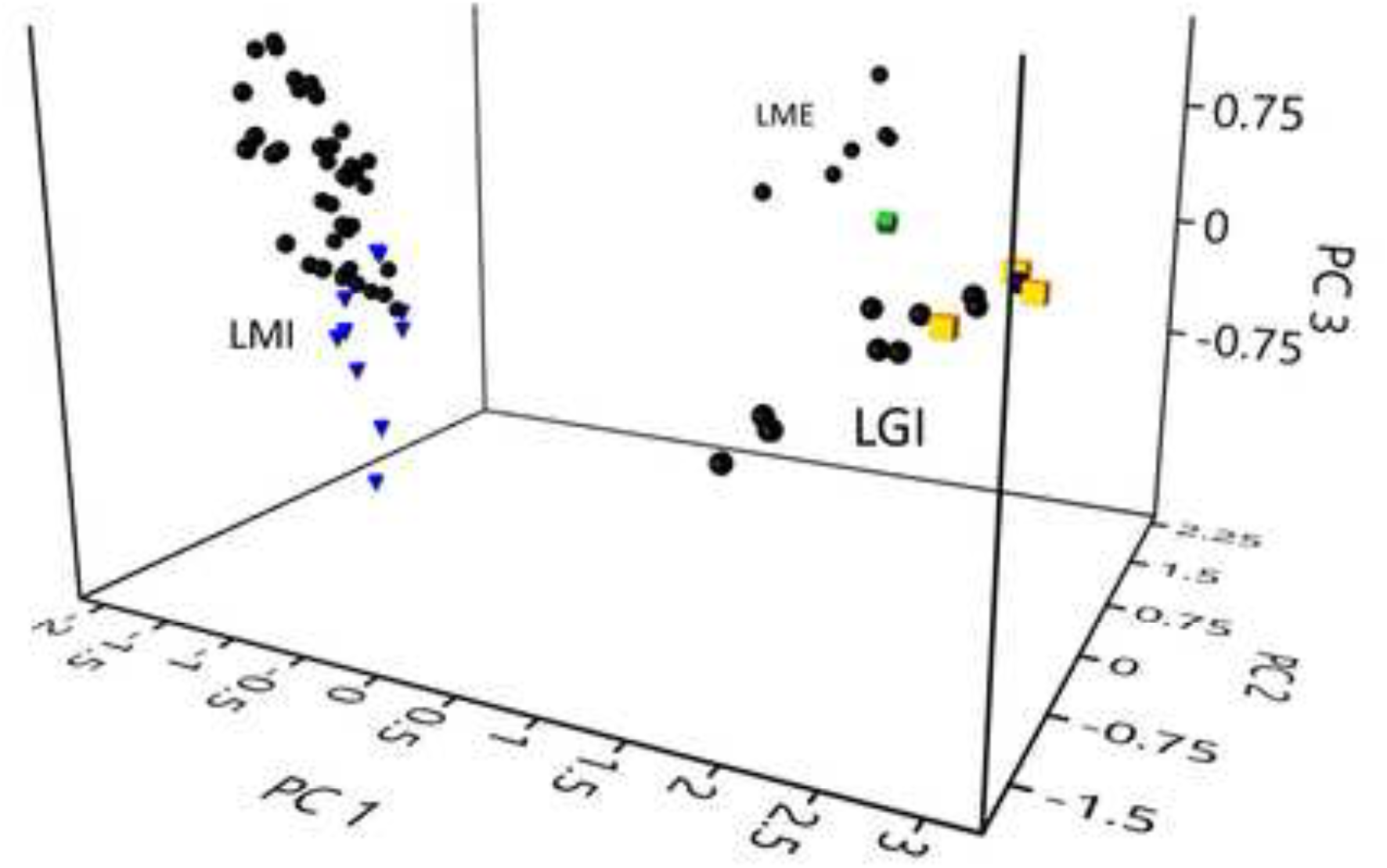
Principal component analysis (PCA) based on pairwise genetic distances calculated from the TBP matrices. Black dots represent clones of the three taxa belonging to the CNR-IBBA duckweed collection. Green diamonds, blue triangles and yellow squares refer to LER-Lme, LER-Lmi and LER-Lgi clones respectively.

## 4. Discussion

This study provides the first direct finding of the interspecific hybrid *L*. × *mediterranea* directly in nature, and not after long-term *in vitro* propagation, unequivocally identified by molecular analysis. Eight specimens, collected at five distinct locations, could not be distinguished from each other either by the nuclear marker TBP or by the *atpF*-*atpH* plastid marker, and are assumed to belong to the same clone LER-Lme. Such triploid hybrid, fully sterile with high probability, as from the lack of seed setting upon flower induction, represents a precious way to trace the spread of purely clonal duckweed lineages over large areas over time. In fact, according to this survey, a single duckweed lineage can spread quite long distance, over a 130 km range, at least. As some of the five collection sites are not directly interconnected by water flow, frond transport through waterbirds by ecto- or endozoochory is the most likely mechanism for their spreading, through a step-by-step process (Coughlan et al., 2017a, Paolacci et al., 2023). Bird-mediated duckweed dispersal over short distances was demonstrated in controlled experimental settings (Coughlan et al., 2017b). By monitoring *L*. *minor* populations in Thuringia (Germany), identical clones of *L*. *minor* were discovered in more than one pond, located at a distance of 1 km and 2.4 km from each other (Bog et al., 2022b). Anthropogenic activities are the second most probable way of propagule dispersal, particularly regarding cross-continental transport of alien species (Fedoniuk et al., 2022, Ziegler et al., 2023). Yearly sampling campaigns including more distant locations could provide interesting data about stability and distribution of such entirely clonal lineages of *L*. × *mediterranea*.

LER-Lme is genetically different from all other *L*. × *mediterranea* clones so far identified within European duckweed germplasm collections (Braglia et al., 2021b; 2024). Four out of seven collection clones were sampled in Italy, at different places and times: clone 9248 was collected in 2000 in the Alpine region Trentino Alto-Adige, clones 9562 and 6861 came from Central Italy, Trasimeno Lake (2016) and Massaciuccoli Lake (1954), respectively, while LM0027 was collected at the Botanical Garden of Naples, at an unknown date, in the same area of LER-Lme. Except for 9248, the Italian clones were found to be genetically distinct homoploid hybrids, having *L*. *minor* as the female parent species, therefore surely independently originated from the LER-Lme clone in this study. The triploid clone 9248, instead, although sharing with LER-Lme the same maternal species, was the most geographically and genetically distant from the other clones in the PCA analysis (Figure 9).

At least five different hybridisation events are then supposed to have originated the distinct clonal lineages so far recovered in Italy, suggesting that recurrent hybridisation of *L*. *minor* and *L*. *gibba*, followed by clone dispersal is not an exceptional occurrence. Accumulation of somatic mutations over time, producing diverging clonal lineages from an original clone cannot be excluded. However, considering the very low mutation rate estimated for *L*. *minor* (Sandler et al., 2020), this would imply very long-term stability of such lineages. Further investigation using higher-resolution markers and population structure analysis will help better understand ancestry and relationships between hybrid clones and their lifespan.

Previous indirect evidence for the presence of the cryptic hybrid in the Campania region comes from the description of sterile *Lemna* specimens resembling *L*. *gibba*, reported as the novel species *L*. *symmeter* (Giuga, 1973). Unfortunately, no specimens from the described populations are known to have been deposited in any herbaria at that time, so we could not evaluate its identity with any other *L.* × *mediterranea* clone. Flower development upon induction by SA in LER-Lme differs from that reported for *L*. *symmeter*, described as characterized by the simultaneous development of the two stamens (from which the name symmeter, meaning symmetric). However, the non-physiological conditions of *in vitro* induction, in contrast with the naturally occurring flowering recorded by Giuga, could be responsible for the observed difference. Large variability of flower development among different clones of both *L. minor* and *L. gibba* was often observed *in vitro* (Landolt, 1980, Fu et al., 2017; Fourounjian et al., 2021). Different maturation of sexual organs, protogyny and homogamy, have been described even within the same species, *L*. *aequinoctialis* Welw., associated with self-sterility or fertility, respectively (Beppu et al., 1985, Lee et al., 2024). A further possibility is that differences among hybrid clones are associated with different genetic makeup, ploidy levels/subgenome composition/kind of cross, between LER-Lme triploid clone and the populations described by Giuga. In accordance with *L. symmeter* description, we can quite surely affirm that LER-Lme is at least self-sterile (possibly fully sterile), as it did not set fruits and seeds under the same induction conditions that were favourable for the parental species. From these and previous data, we then conclude that *L.* × *mediterranea* matches the description of *L*. *symmeter* and that the Italian peninsula represents an area favourable to *L.* × *mediterranea* origin and propagation. Further field studies may reveal additional hotspots for *L. gibba* and *L. minor* hybridisation. To this regard, promising sites could be Northern Germany and the Netherland, where morphologically intermediate forms between *L. minor* and *L. gibba* were described in the past (De Lange and Pieterse, 1973, Landolt, 1975).

The two parental species *L*. *minor* and *L*. *gibba* were both largely present in the study area but, likely because of different ecological preference (Landolt, 1987), never co-occurred in the same waterbody, suggesting this is not the most common occurrence. In agreement with recent reports (Bog et al., 2022b, Senevirathna et al., 2023, Schmid et al., 2024) *L*. *minor* was associated with high intraspecific genetic variability, scorable by the high polymorphic TBP marker: out of thirteen specimens, 10 have distinct allelic patterns. However, among all the isolated accessions of *L*. *gibba* and *L*. *minor* we could find just two candidates as the putative parental clones, LER016 (*L*. *gibba*) and LER017 (*L*. *minor*) that showed the same allele combinations at β-tubulin loci found in LER-Lme. Evidence for recurrent hybridisation led us to suppose that flowering in *L*. *minor* is more frequent than often reported. Then, cross-pollination between the two species, when fronds are in close proximity, although unlikely, should give fertile seeds. Hybrids may locally have some competitive advantage over parent species allowing them to propagate and colonize new water bodies. Cross-pollination experiments are now ongoing to reproduce and test the success rate of interspecific hybridisation *in vitro*.

Another evidence in favour of underestimated crossing rates in the parent species comes from the high intraspecific variability highlighted in this study. Both parent species, particularly *L*. *minor*, showed intraspecific diversity between populations (sampling sites), as estimated by TBP. This agrees with recent population studies on *L*. *minor* worldwide, using different markers. Bog et al. (2022b) by using AFLP, found 20 distinct clones based on 36 samples collected in a small area in Thuringia, in Germany, some of which living side by side in the same pond; Senevirathna et al. (2023) identified at least three distinct genetic clusters among 30 samples of *L*. *minor* from eight sites in Alberta (Canada) by using Genotyping By Sequencing, with sampling sites containing individual from the three clusters; by the same approach, Schmid et al. (2024) identified high inter-population diversity between 23 sampling sites across Switzerland, represented by eight distinct lineages. In accordance, in this study we showed that *L. minor* populations can be polyclonal and that individual clones can be found at different sites over large distances.

The potential success of interspecific *Lemna* hybrids is clearly witnessed by *L*. *japonica* Landolt, described by E. Landolt as a species distinct from *L*. *minor* in 1980, native to East Asia (Landolt, 1980). Recent analysis revealed that this species is an interspecific hybrid between *L*. *minor* and *L*. *turionifera* Landolt (Braglia et al., 2021b, Ernst et al., 2023, preprint). In the wide original area, extending from Russia to China and including Japan and Korea, the two parent species are indeed sympatric. However, molecular analysis revealed that a large number of clones distributed across Asia and Europe and classified as *L*. *minor* belong instead to this cryptic taxon, despite the absence of the parent species *L*. *turionifera* in most of these areas (Braglia et al., 2021b, Volkova et al., 2023). Schmid et al. (2024) recently reported that *L*. *japonica* is widespread throughout Switzerland, by far distant from its putative centre of origin. As *L*. *japonica* has never been seen to set seeds, it is supposedly sterile as most interspecific hybrids. Its migration history must solely rely on asexual propagation and long-distance dispersal over time. Even in this case, recurrent hybridisation is likely, as both homoploid and triploid clones with different subgenome composition have been found among collection accessions (Ernst et al., 2023, preprint; Michael T., unpublished results).

Other species found across the study area in Italy included *W*. *arrhiza*, *L*. *trisulca* (at one site only) and the alien species *L*. *minuta*. While *L*. *minor* and *L*. *gibba* are both autochthonous species commonly reported in Italy (Pignatti et al., 2017, Landolt, 1986), *L*. *minuta* is an alien species, native to the American continent, but now largely distributed in Europe and the Italian peninsula where it has been reported since 1989 (Desfayes, 1992). Since then, *L*. *minuta* has been widely reported in Northern and Central Italy, where it is described as naturalized or invasive in different regions (Iamonico et al., 2010, 2012; Ceschin et al., 2016), in competition with *L*. *minor*. In Southern Italy, *L*. *minuta* stations were found in Apulia (Beccarisi at al., 2006), Sicily (Marrone and Naselli-Flores, 2011) and Calabria (Salerno and Ceschin 2015). The only record of *L*. *minuta* in Campania dates to 2017 and reported the species as casual (Stinca et al., 2017). The present data suggest that *L*. *minuta,* although spreading from North to South, did not have the same character of invasiveness as in northern regions. The possibility of a stronger competition by *L*. *gibba* and, possibly, *L*. × *mediterranea*, more common in the warmer southern regions than in northern ones, is worth investigating.

No alien *Wolffia* species were reported in this survey. Although *W. arrhiza*, distributed in Europe and Africa, is the only autochthonous *Wolffia* species in Italy, other alien species are in fact spreading in Europe from other regions. *W*. *columbiana* H. Karst. has been described in Europe (Dieter et al., 2020) and more recently also in Italy (Ardenghi et al., 2017) and *W. globosa* (Roxb.) Hartog & Plas has been expanding from Asia to Europe since 2010 (Kirjakov and Velichkova 2013).

No difference was observed in this study with respect to the preference of any duckweed species for the two main kind of investigated waterbodies, NW, chiefly different traits of the medium size Volturno river and AW, mainly slow flowing canals.

Additional sampling campaigns, supported by molecular analysis, may reveal unknown aspects of *L*. × *mediterranea* distribution and the bases for their success.

**Supplementary Table 1.** List of the duckweed clones belonging to the CNR-IBBA dataset collection included in the present paper. The *ex-situ* germplasm repository is also available at the present link - https://scientific-collections.gbif.org/collection/24fdf8b9-2165-44a5-9e30-019a2190331d

**Supplementary Table 2.** Dissimilarity values collected by a pairwise comparison analysis based on Whittaker’s index formula.

**Supplementary Figure 1.**
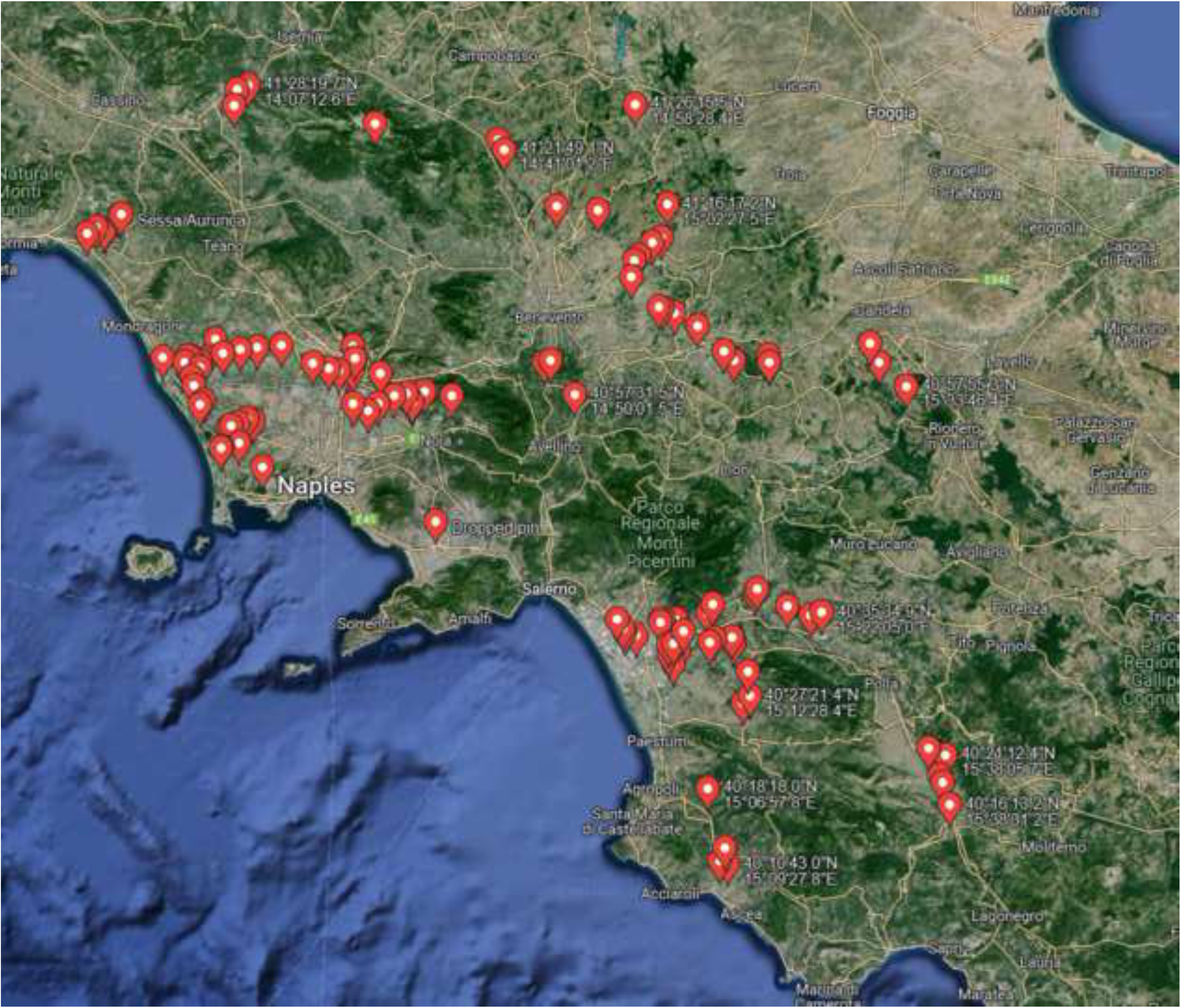
Map of the sampling campaign. Red pins represent the places visited during the field expedition. Red circles represent the areas previously described by Giuseppe Giuga (1973), in which he sampled a putative species named as *L. symmeter,* never formally described.

**Supplementary Figure 2.**
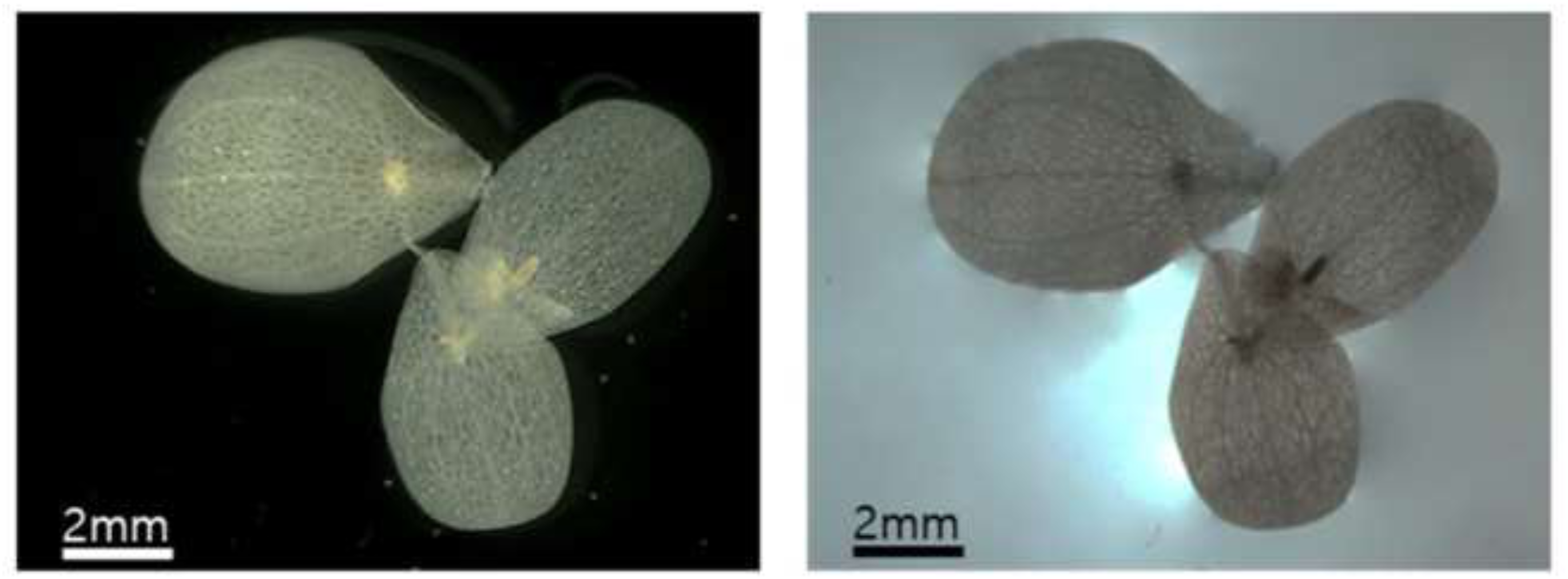
Representative stereomicroscope images of cleared fronds colonies for the determination of vein number per frond in the accession LER021 (*L.* × *mediterranea*). Fronds colonies were observed under bright- and dark-field conditions. Bar = 2mm.

**Supplementary Figure 3.**
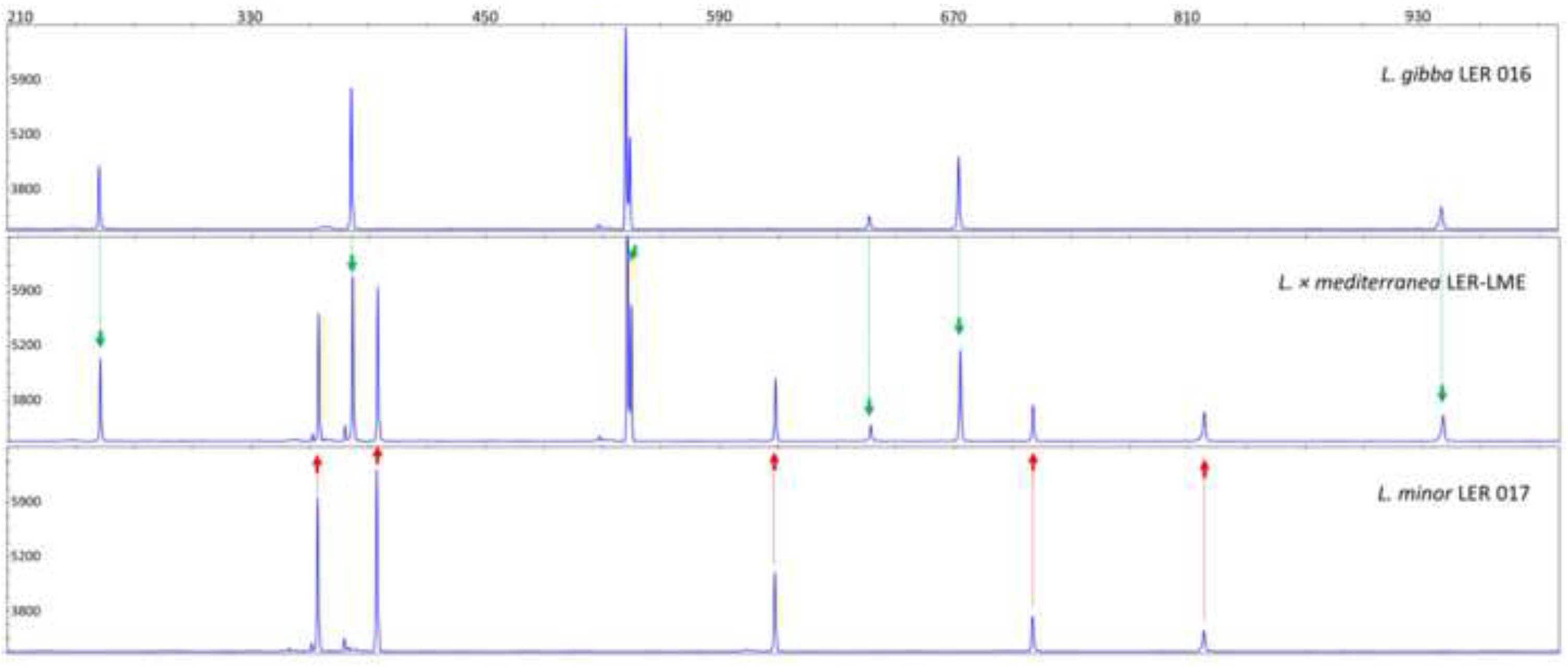
Comparison among Capillary Electrophoresis TBP peak profiles of a *L.* × *mediterranea* LER-Lme, and the putative parental clones *L. minor* LER017 and *L. gibba* LER016. Traceable alleles (peaks) from parents to the hybrid are indicated by green and red arrows referring to *Lemna gibba* and *L. minor* respectively.

## Acknowledgments

We are indebted to Dr. Jörg Fuchs, from The Leibniz Institute of Plant Genetics and Crop Plant Research (IPK), Gatersleben, DE, for providing data on absolute genome size measurements of clone LER0027, 9248 and 9425a. The authors want to thank Dr. Luigi Gennaro Izzo and Dr. Maurizio Iovane, University of Naples, for their immense contributions to sampling the Lemnaceae plants. Special thanks go to Prof. Klaus J Appenroth, University of Jena (DE) for critical reading of the manuscript.

## Author Contributions

Conceptualization, L.M., L.E.R.; methodology, L.E.R., L.B., L.M.; formal analysis, L.B., L.M., L. E.R.; investigation L.E.R., Y.L., M.A.I, L.B., F.G, S.G.; data curation L.B; L. E.R.; writing - original draft, L.M., L.E.R.; writing - review and editing, L.B., L.M., M.A.I, L. E.R., Y.L., S.G, F.G.; supervision, L.M., L.B.; funding acquisition, L.E.R, L.B., L.M. All authors have read and agreed to the published version of the manuscript.

## Data Availability Statement

Plastid barcoding sequences will be published in a public sequence repository (NCBI).

